# MultiNMRFit: A software to fit 1D and pseudo-2D NMR spectra

**DOI:** 10.1101/2024.12.19.629408

**Authors:** Pierre Millard, Loïc Le Grégam, Svetlana Dubiley, Valeria Gabrielli, Thomas Gosselin-Monplaisir, Guy Lippens, Cyril Charlier

## Abstract

Nuclear Magnetic Resonance (NMR) is widely used for quantitative analysis of metabolic systems. Accurate extraction of NMR signal parameters – such as chemical shift, intensity, coupling constants, and linewidth – is essential for obtaining information on the structure, concentration, and isotopic composition of metabolites. We present MultiNMRFit, an open-source software designed for high-throughput analysis of one-dimensional NMR spectra, whether acquired individually or as pseudo-2D experiments. MultiNMRFit extracts signal parameters (e.g. intensity, area, chemical shift, and coupling constants) by fitting the experimental spectra using built-in or user-defined signal models that account for multiplicity, providing high flexibility along with robust and reproducible results. The software is accessible both as a Python library and via a graphical user interface, enabling intuitive use by end-users with or without computational expertise. We demonstrate the robustness and flexibility of MultiNMRFit on ^1^H, ^13^C, and ^31^P NMR datasets collected in metabolomics and isotope labeling studies.

**Availability and Implementation:** MultiNMRFit is implemented in Python 3 and was tested on Unix, Windows, and MacOS platforms. The source code and the documentation are freely distributed under GPL3 license at https://github.com/NMRTeamTBI/MultiNMRFit/.

**Supplementary data:** Supplementary data are available online.

**Contact:** Cyril Charlier and Pierre Millard

## 1. Introduction

Nuclear Magnetic Resonance (NMR) spectroscopy is a powerful tool for investigating metabolism. Key applications include identifying and quantifying metabolites in metabolomics, determining the isotopic content of metabolites in ^13^C-fluxomics, and identifying and characterizing protein-metabolite interactions (Koczula *et al*., 2016; Moco, 2022; Nargund *et al*., 2013; Diether *et al*., 2019). Extracting meaningful information from NMR spectra requires precise characterization of each signal of interest, including their position (chemical shift), intensity (and area) and shape (coupling constants and linewidth) (Edison *et al*., 2021). Signal parameters can be estimated by fitting spectra using various mathematical models that relate these parameters to the observed signals. This fitting step is often achieved with in-house scripts, which, while functional and flexible, may lack the robustness and reproducibility necessary for large-scale or complex NMR studies of metabolic systems. Commercial software, such as TopSpin (Bruker), Chenomx (Chenomx Inc), and Mnova (Mestrelab), offer user-friendly alternatives, but function as proprietary “black boxes” and are limited to predefined signal models. To enhance accessibility and reproducibility, open-source libraries for R, MATLAB, and Python have been developed, including MetaboLab (Ludwig and Günther, 2011), MetaboDecon1D (Häckl *et al*., 2021), BATMAN (Hao *et al*., 2012, 2014), AQuA (Röhnisch *et al*., 2021), and FitNMR (Dudley *et al*., 2020). However, they are designed for use via scripting and often lack features essential for users without computational expertise. Such missing features include statistical methods for calculating confidence intervals of extracted parameters, graphical user interfaces, and logging systems to ensure reproducibility by saving calculation parameters and process information along with the results. Another key challenge is that most existing tools treat each peak as an independent signal (Smith, 2017). However, the NMR signal of a single nucleus often consists of multiple peaks arising from homo- and heteronuclear scalar couplings, which are intrinsic properties crucial for identifying metabolites and quantifying their isotopic content. While tools like Dmfit for solid and liquid NMR data (Massiot *et al*., 2002) and, more recently, the R package rnmrfit (Sokolenko *et al*., 2019) account for multiplicity, most current software does not, which limits their effectiveness in extracting and utilizing this information.

Consequently, detailed analysis of NMR spectra remains a low throughput task, typically requiring expert users, which can limit its scalability for large datasets such as those generated in real-time NMR experiments. The development of user-friendly, open-source software that incorporates signal multiplicity and supports flexible implementation of tailored signal models would significantly enhance the scope, efficiency, quality, and reproducibility of NMR-based metabolic studies.

To address these challenges, we have developed MultiNMRFit, a versatile Python software for fitting one dimensional NMR spectra, whether acquired individually or as pseudo-2D datasets in time-course experiments. MultiNMRFit is generic and can be applied to any nucleus (as shown here for ^1^H, ^13^C, and ^31^P spectra), any type of 1D or pseudo-2D NMR experiment, and any type of signal. It includes built-in signal models for common multiplets and is easily extendable with user-defined models to process less conventional signals. MultiNMRFit is accessible both as a Python library and via a graphical user interface. This software is intended for both expert and non-expert NMR users, regardless of their computational expertise. We demonstrate the robustness of MultiNMRFit on synthetic data and through its application to real-time metabolomics and isotope labeling experiments.

## 2. Methods and implementation

The general workflow of MultiNMRFit is shown in Figure 1. MultiNMRFit is a Python-based library that operates within common web browsers via a Streamlit-powered graphical user interface. Users can load 1D and pseudo-2D Bruker spectra pre-processed beforehand (*i*.*e*. referenced, phase-corrected and baseline-corrected). Spectra can also be provided as text files containing chemical shift values and corresponding intensities. MultiNMRFit performs automated peak picking within a specified spectral region, and users can add additional peaks manually. One or several peaks can then be grouped into a signal, and a model must be selected for each signal. As detailed in the next section, each model contains equations to simulate the corresponding signal based on parameters. MultiNMRFit automatically suggests a list of models based on the number of peaks (e.g., *doublet of doublet*s and *quartet* for signals containing four peaks). Note that a given peak can be included in multiple signals to account for overlapping peaks. The initial values and bounds of each signal’s parameters are automatically estimated from the peak list to ensure reliable convergence and can be refined by users if needed. For example, the initial chemical shift and intensity are taken directly from the peak list, while coupling constants are estimated from the difference in chemical shifts between peaks. Bounds can be defined as either absolute or relative values, and their default values (contained in the model files) can be adjusted by users to better reflect their experimental spectra. For spectrum fitting, parameters *p* are estimated by minimizing the cost function *c*(*p*), which is defined as the sum of squared residuals between the experimental intensities (*a*_*i*_) and the values simulated by the model *y*_*i*_(*p*):

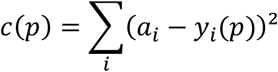

**Figure 1.**
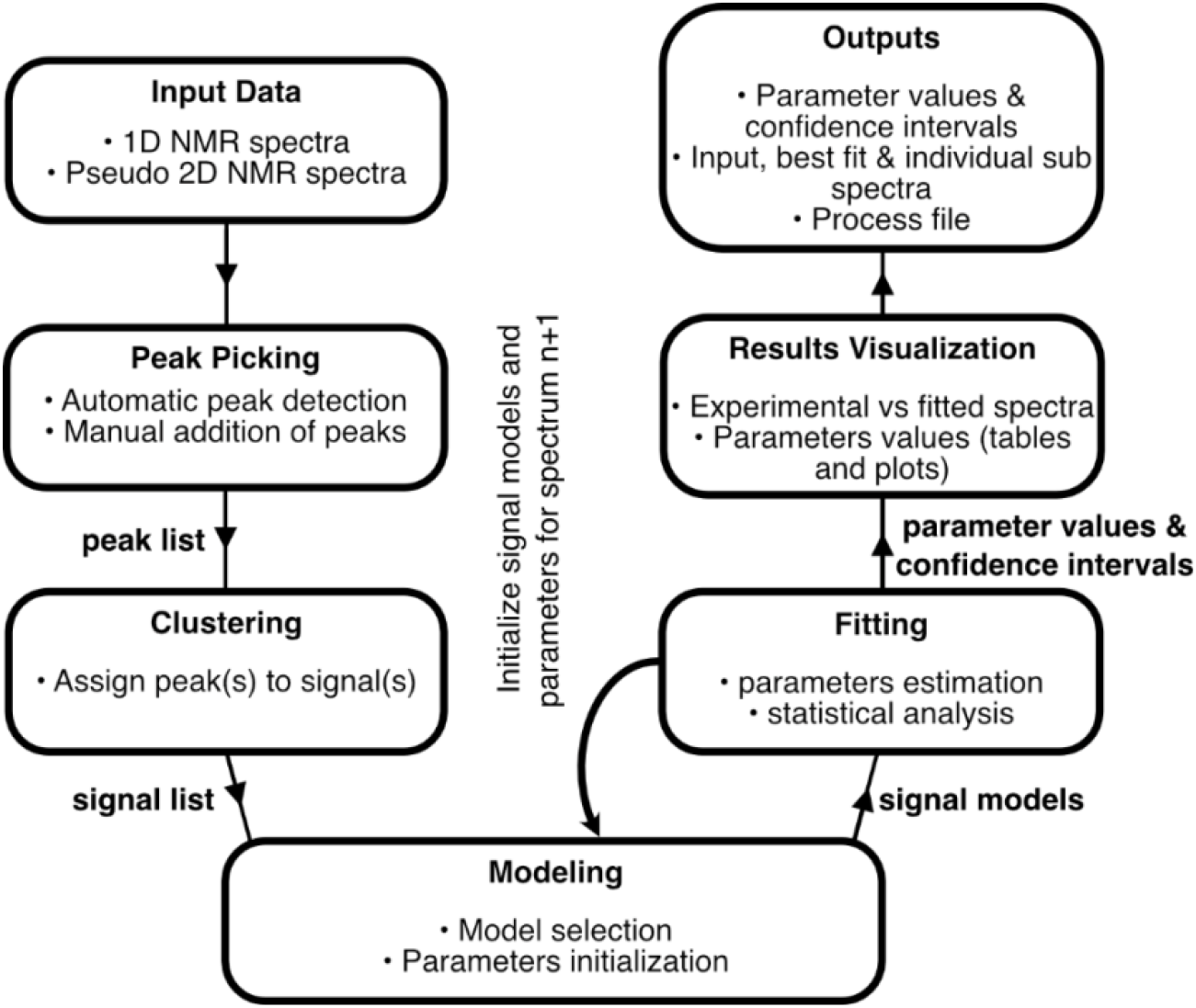
Flow chart of MultiNMRFit.

The cost function is minimized using the L-BFGS-B method (Byrd *et al*., 1995), with an optional initial refinement step performed by the differential evolution algorithm (Storn and Price, 1997). Standard deviations on parameters are estimated from the covariance matrix. For batch processing of spectra, users can select a spectrum as reference to automatically initialize signals and parameters values for the next spectra, thereby automating the processing of multiple spectra for high-throughput analysis. Finally, MultiNMRFit provides a range of interactive plots for visual inspection of fitting results. The results can be exported as *csv* files containing *(i)* parameter values (with their confidence intervals) and *(ii)* chemical shifts and intensities of experimental and fitted spectra.

The complete process is automatically saved as a pickle file, which can be reloaded in MultiNMRFit to continue previous work or reanalyze calculation results. This file can also be shared, enabling collaborative analysis across multiple users and ensuring reproducibility. Detailed usage of MultiNMRFit can be found in the online documentation (https://multinmrfit.readthedocs.io).

## 3. Signal models

A signal model consists of *(i)* equations that describe signal intensity as a function of chemical shift (used for simulations) and *(ii)* a list of parameters with their default values and bounds (used for optimization). Various mathematical functions, including Lorentzian, Gaussian, and Voigt functions, can be used to simulate the NMR spectrum of a given signal. For example, the intensity *y*_*i*_ of a doublet can be described using the following mixed Gaussian-Lorentzian function:

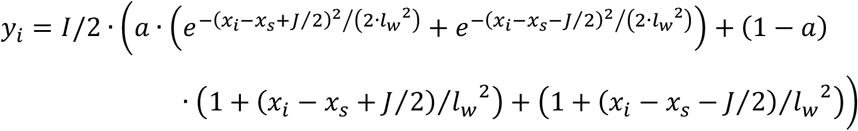

with parameters *x*_*s*_ (chemical shift of the signal), (total signal intensity), (scalar coupling constant), (linewidth at half maximum), and *a* (fraction of Gaussian part of the signal). An NMR spectrum containing multiple signals can be simulated by summing the spectra of individual signals.

By default, MultiNMRFit provides mixed Gaussian-Lorentzian models for common multiplets, including singlet, doublet, triplet, doublet of doublets, and quadruplet. Advanced users can create custom models to handle less conventional signals, such as signals with other coupling patterns or described using alternative mathematical functions. A model template is included with MultiNMRFit for this purpose, and a detailed tutorial on model construction is available in the documentation.

## 4. Validation with synthetic datasets

We validated MultiNMRFit using synthetic datasets (Supplementary data). Briefly, we generated spectra containing two signals (a triplet and a quadruplet) with varying degrees of overlap and noise, and we used MultiNMRFit to retrieve the parameters of each signal. Results confirm the capability of the software to handle complex spectral data and deliver robust parameter estimation even under challenging conditions (such as low signal-to-noise ratio or significant signal overlaps, Supplementary Figures S1 and S2). Moreover, fitting multiplets rather than individual peaks increased the accuracy and precision of the estimated parameters (Supplementary Figure S3).

## 5. Application to metabolomics datasets

We used MultiNMRFit to monitor the dynamic conversion of glucose through the initial steps of glycolysis. Glucose was added to a buffer solution containing sodium trimethylsilyl propionate (TSP, used as internal standard), ATP and the first three enzymes of glycolysis: glucokinase, glucose-6-phosphate isomerase, and phosphofructokinase (Figure 2A, Supplementary data). The time-course signals of glucose (GLC), glucose-6-phosphate (G6P), fructose-6-phosphate (F6P), fructose-1,6-bisphosphate (FBP), ATP and ADP were monitored in real-time using a pseudo-2D experiment consisting of 1024 1D ^1^H spectra. All signal parameters were successfully extracted, including for overlapping signals of ADP and ATP (Figure 2B). The integrals of signals specific to each metabolite were normalized to the TSP signal (Figure 2C). A progressive decrease in GLC signal was observed, accompanied by a transient accumulation of G6P and F6P, which were eventually converted into FBP. The high precision and resolution on the estimated dynamics illustrate the value of MultiNMRFit for high-throughput processing of large datasets, with the relative concentrations of the six metabolites of interest quantified in less than three hours for the 1024 spectra. Simultaneously the ^31^P spectra were acquired along the way (Cox *et al*., 2019) and were equally fitted using MultiNMRFit (Figure S4), To demonstrate the applicability of MultiNMRFit for other nuclei, we fitted ^13^C NMR spectra obtained from a separate experiment in which we monitored the conversion of uniformly ^13^C-labeled GLC to G6P by hexokinase using a ^13^C detected pseudo-2D experiment (Supplementary data). In this experiment, both GLC and G6P give rise to a doublet of triplets. Here again, the spectra could be fitted accurately, and the concentration dynamics of the two ^13^C-labeled reactants could be obtained from real-time data (Supplementary Figure S5).

**Figure 2.**
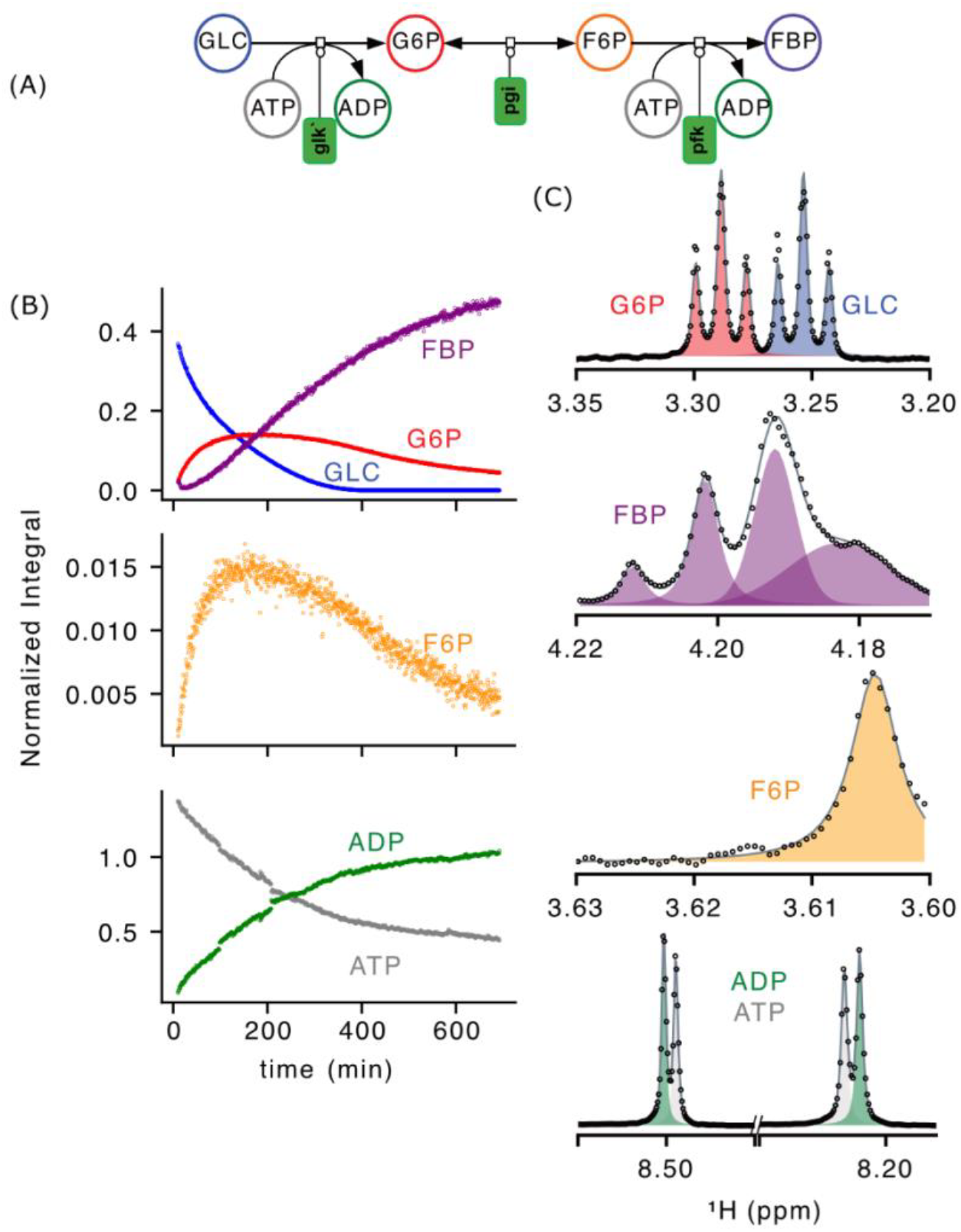
Applications of MultiNMRFit to multi-enzymatic reactions. **(A)** Scheme of the biochemical pathway monitored in a real-time NMR experiment. **(B)** Dynamic of integrals of glucose (GLC, blue), glucose-6-phosphate (G6P, red), fructose-6-phosphate (F6P, orange), fructose-1,6-bisphosphate (FBP, purple), adenosine triphosphate (ATP, gray), and adenosine diphosphate (ADP, green). Integrals were normalized with the TSP signal. **(C)** Examples of signals processed with MultiNMRFit for each molecule of interest, where individual signals are shown by the shaded areas and the experimental and fitted spectra are represented by dots and lines, respectively.

## 6. Application to isotopic studies of metabolism

MultiNMRFit can also exploit the multiplicity resulting from the presence of isotopes in metabolites to quantify their isotopic content – a key capability for applications such as detecting counterfeit chemicals, identifying metabolic pathways, and quantifying metabolic fluxes (Ribeiro *et al*., 2018; Christensen *et al*., 2002).

To demonstrate this, we conducted a ^13^C-isotope labeling experiment by growing *Escherichia coli* on a mixture of glucose (15 mM, 80% 1-^13^C_1_-glucose + 20% U-^13^C_6_-glucose) and acetate (8 mM, 100% 2-^13^C_1_-acetate). Samples were collected each hour to quantify the dynamics of all isotopic forms of acetate (Supplementary data). In the ^1^H NMR spectra, the methyl protons of acetate may give rise to nine peaks corresponding to the four isotopic species of this compound: a singlet for ^12^C_2_-acetate, a doublet with a small (6 Hz) ^2^*J* coupling constant for 1-^13^C_1_-acetate, a doublet with a large (127 Hz) ^1^*J* coupling constant for 2-^13^C_1_ acetate, and a doublet of doublet for U-^13^C_2_-acetate (Figure 3 and S6). The results showed that the total acetate concentration remained stable throughout the experiment (8.5±0.1 mM), which could be interpreted as an acetate flux close to zero (Figure 3C). However, while the provided isotopic form (2-^13^C_1_-acetate) initially represents 99% of the total acetate pool, its concentration decreased over time (from 8.5 to 6.5 mM). Concurrently, the concentration of the other labeled forms (derived from glucose) increased (from 0.1 to 1.9 mM). These results support previous findings that acetate is continuously exchanged between cells and their environment, even in the absence of net acetate accumulation in the medium (Enjalbert *et al*., 2017).

**Figure 3.**
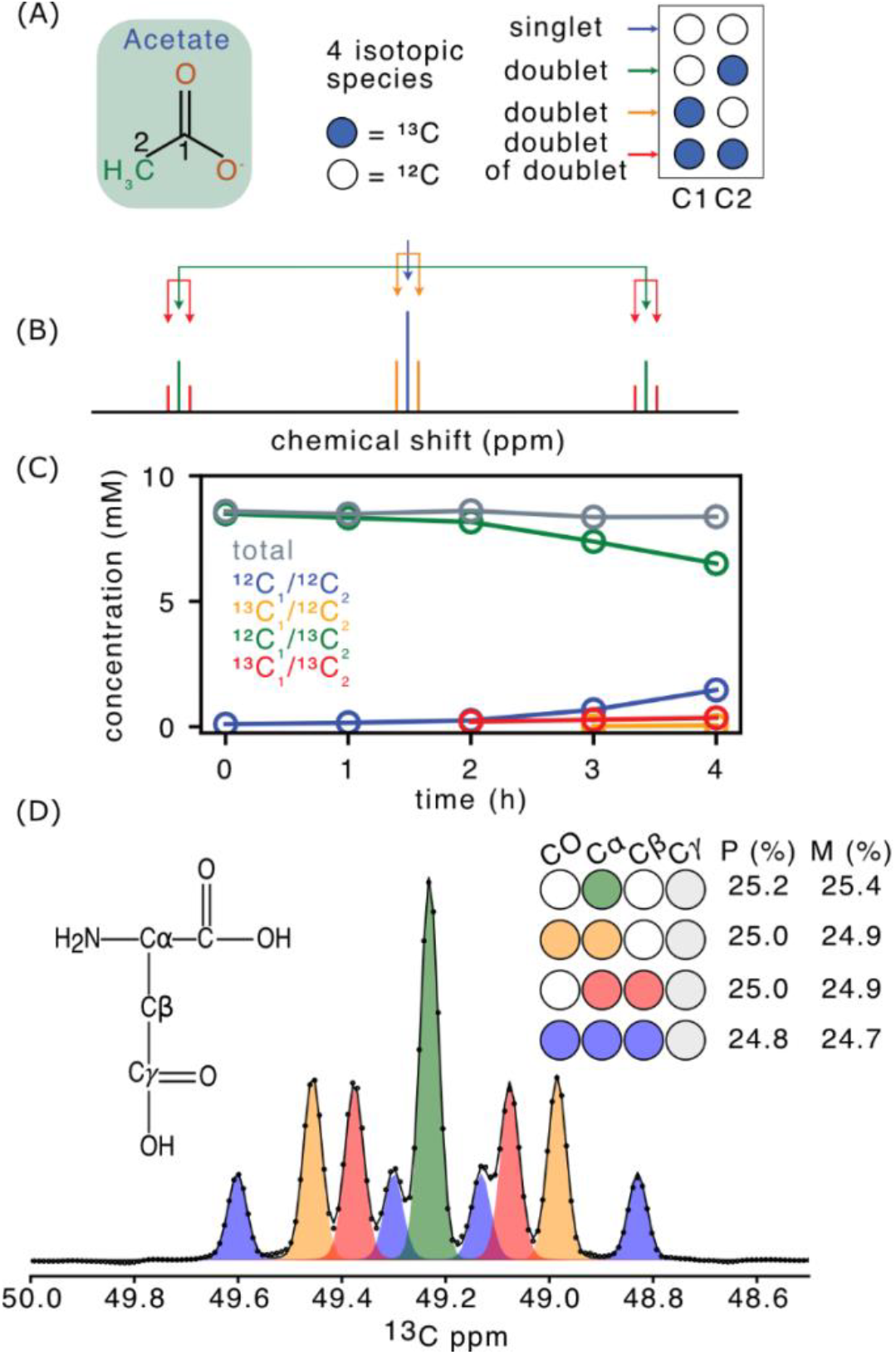
Applications of MultiNMRFit to isotope labeling studies. **(A)** Isotopic forms of acetate. **(B)** Theoretical ^1^H spectrum of the methyl protons of acetate, with a singlet for ^12^C_2_-acetate (blue), a doublet for 1-^13^C_1_-acetate (orange), a doublet for 2-^13^C_1_ acetate (green), and a doublet of doublets for U-^13^C_2_-acetate (red). **(C)** Time-course concentrations of the four isotopic forms of acetate (using the color code shown in panel B) and of the total acetate concentration (in grey). **(D)** Experimental spectrum of ^13^C-labeled aspartate focused on the Cα signal processed with MultiNMRFit, with the experimental shown as dots, the fitted spectrum as a line, and individual signals as shaded areas. On the right side the different labeling scheme are displayed. The table corresponds to the predicted (P) and the measured (M) fractions of each isotopic species.

As an additional test case, we performed a ^13^C-labeling experiment to generate a standard sample with known and predictable labeling patterns of metabolites, which enables validation of analytical and processing methods (Millard *et al*., 2014). We analyzed the labeling pattern of aspartate using a ^1^H-^13^C HSQC experiment, which provides information on four isotopic species (Figure 3D). Quantification results were in close agreement with the predicted labeling pattern of aspartate (differences below 1%), demonstrating that MultiNMRFit can accurately quantify the isotopic content of metabolites from ^13^C-NMR spectra.

These examples illustrate the applicability of MultiNMRFit for extracting accurate quantitative isotopic information on metabolites from ^1^H and ^13^C NMR spectra.

## Conclusion

We present MultiNMRFit, an open-source Python software designed to extract detailed quantitative parameters from NMR spectra. MultiNMRFit is applicable to spectra acquired from any nucleus, whether as 1D or pseudo-2D spectra, and can handle any type of multiplicity through a flexible system for implementing NMR signal models. MultiNMRFit is accessible to non-expert users through a graphical user interface and can also be used as a Python package, enabling seamless integration into metabolomics and fluxomics workflows. Furthermore, MultiNMRFit holds significant potential for other NMR applications, including protein studies, the pharmaceutical industry, biotechnology, and chemistry.

## Acknowledgements

We thank Marin Martin Mawah and Pauline Rouan for testing previous versions of the software and for insightful discussions. We thank Dr. Edern Cahoreau and Lindsay Peyriga (MetaToul, Toulouse metabolomics & fluxomics facilities, www.metatoul.fr) for access to NMR facilities. MetaToul is part of MetaboHUB, the French National Infrastructure for Metabolomics and Fluxomics (www.metabohub.fr).

## Author contributions

**PM & CC** Software, Project administration, Supervision, Conceptualization, Writing - review & editing; **LL** Software**; SD, VG & TM** Resources; **GL** Conceptualization, Writing - review & editing.

