## Supplementary material for "MultiNMRFit: A software to fit 1D and pseudo-2D NMR spectra"

1. VALIDATION OF MULTINMRFIT ON SYNTHETIC DATASETS 3

1.1. ANALYSIS OF ISOLATED AND OVERLAPPED SIGNALS 3
1.2. COMPARISON OF MULTIPLETS VS SINGLETS 4

2. APPLICATION ON METABOLOMICS AND ISOTOPIC DATASETS 5

2.1. SAMPLE PREPARATION 5

2.1.1. EXPRESSION AND PREPARATION OF ENZYMES FOR REAL-TIME MONITORING OF BIOCHEMICAL REACTIONS 5
2.2.2. CULTIVATION AND SAMPLE PREPARATION FOR $^{13}C$-ISOTOPE LABELING EXPERIMENTS 5

2.2. NMR SPECTROSCOPY AND DATA PROCESSING 6

2.2.1. REAL-TIME ANALYSIS OF BIOCHEMICAL REACTIONS BY $^{1}H$ AND $^{31}P$ NMR 6
2.2.2. REAL-TIME ANALYSIS OF BIOCHEMICAL REACTIONS BY $^{13}C$ NMR 7
2.2.2. DETERMINATION OF $^{13}C$-ISOTOPE INCORPORATION IN ACETATE 8
2.2.2. DETERMINATION OF $^{13}C$-ISOTOPE INCORPORATION IN ASPARTATE 9

3. REFERENCES 9

**Data availability:** All data and code required to reproduce the analyses and figures provided in this Supplementary material are available as a Zenodo repository at: https://doi.org/10.5281/zenodo.14528296

### 1. Validation of MultiNMRFit on synthetic datasets

The robustness of MultiNMRFit was evaluated using synthetic datasets. Accuracy was calculated as the (relative) mean error between estimated and true parameter values, and precision was estimated as the (relative) standard deviation on the estimated parameters:

$$accuracy = \frac{|p_{true} - p_{estimated}|}{p_{true}}$$

$$precision = \frac{standard\ deviation(p_{estimated})}{p_{true}}$$

#### 1.1 Analysis of isolated and overlapped signals

First, we simulated 4 sets of spectra, each containing a triplet with a signal-to-noise ratio (S/N) ranging from 2.5 to 20. For each S/N, 250 spectra were generated with random noise, and MultiNMRFit was used to estimate the accuracy and precision of each estimated parameter (chemical shift, coupling constant, linewidth, and intensity). The accuracy and precision of all parameters (Figure S1) remained below 5% and 15%, respectively, demonstrating that MultiNMRFit provides robust parameter estimation on an isolated signal, even at low S/N.

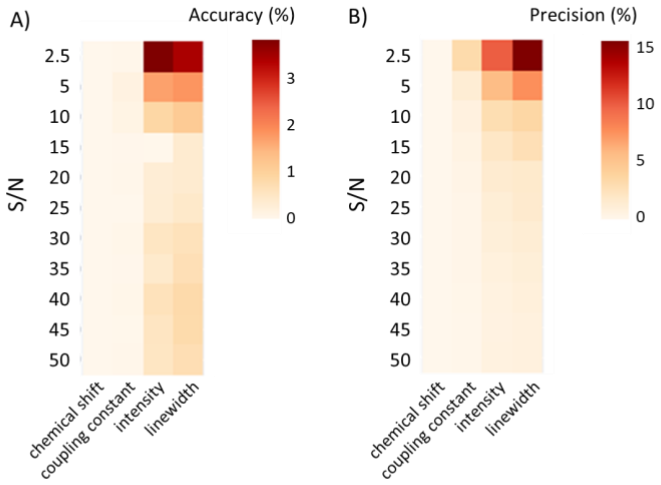


**Figure S1. Validation of MultiNMRFit on an isolated signal**. Evaluation of the robustness of MultiNMRFit on spectra containing a triplet at various signal-to-noise (S/N) ratio. For each S/N ratio, 100 spectra were simulated with random noise, and MultiNMRFit was used to estimate all parameters, from which the accuracy and precision were determined. Simulations were carried out using the triplet model of MultiNMRFit with the following input: magnetic field ($B_0$) of 800 MHz, a chemical shift at 7.5 ppm, a coupling constant (J) of 7 Hz, a linewidth (lw) of 1 Hz, a gaussian/lorentzian ratio of 0.5, and a total intensity of $4.10^5$. Spectra were simulated with 1024 points spanning a 1 ppm region. The signal-to-noise was determined on the down-field satellite of the triplet.

Second, we applied the same analysis to a more complex spectrum containing two signals (a triplet and a quadruplet) with varying degrees of overlap (separated by 30, 20 or 15 Hz), across a broad range of S/N (Figure S2). Again, all spectra were fitted satisfactorily (examples are shown in Figure S2C), and all parameters were estimated for both multiplets with accuracy and precision below 10% (Figure S2A-B), even in cases of significant signal overlap. This highlights the capability of MultiNMRFit to deliver robust parameter estimation for spectra with overlapped signals, even at low S/N.

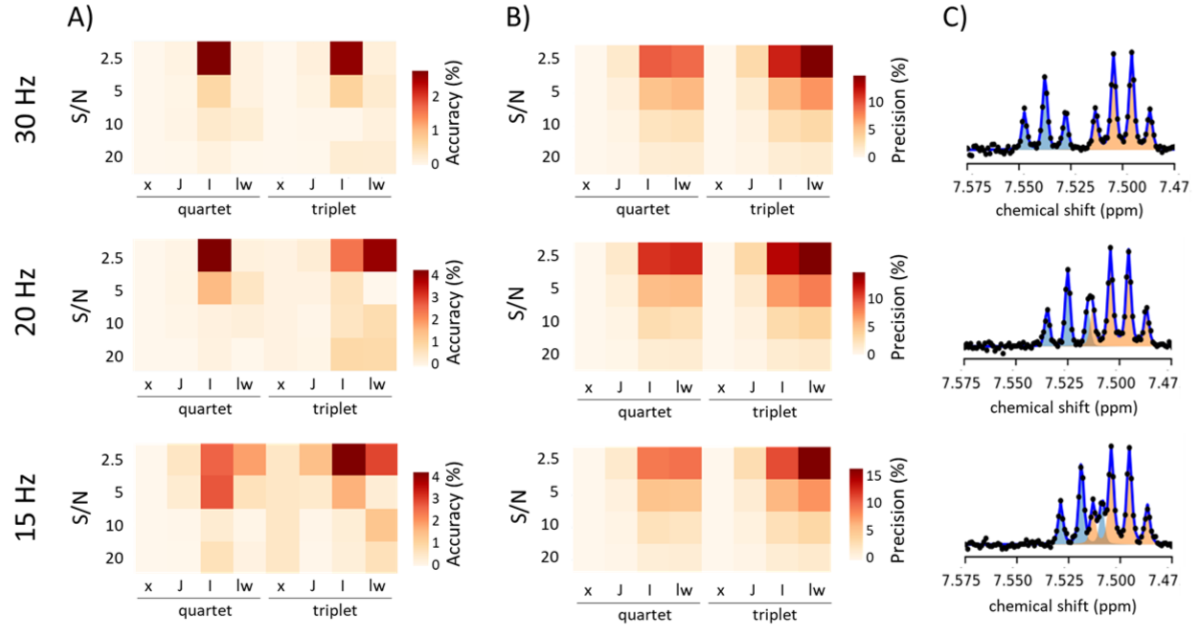


**Figure S2. Validation of MultiNMRFit on overlapped signals**. Accuracy **(A)** and precision **(B)** of all parameters (x: chemical shift, J: coupling constant, I: intensity, lw: linewidth) obtained by MultiNMRFit for a spectrum containing a triplet and a quadruplet with different degrees of overlap (separation of 15, 20 or 30 Hz between the center of both multiplets) and different S/N ratio. Simulations were carried out using the triplet and quadruplet models of MultiNMRFit with the following input: magnetic field ($B_0$) of 800 MHz, a chemical shift at 7.5 ppm, a coupling constant (J) of 7 Hz, a linewidth (lw) of 1 Hz, a gaussian/lorentzian ratio of 0.5 and a total intensity of $4.10^5$. Spectra were simulated with 1024 points spanning a 1 ppm region. The signal-to-noise was determined on the down-field satellite of the triplet. Examples of spectra are shown in panel **(C)**, where the dots represent the simulated spectra, the blue line represents the best fit obtained by MultiNMRFit, and the blue and orange areas represent the individual signals.

#### 1.2 Comparison of multiplets vs several singlets

Most NMR processing tools treat each peak as an independent signal and rely solely on singlets in the fitting process. We compared the accuracy and precision of parameter estimates obtained using either singlets or multiplets. It is important to note that, when using singlets, certain parameters - such as coupling constants - cannot be directly extracted from the fitting process. Instead, users must recalculate them based on the estimated chemical shifts of individual singlets. Therefore, for the purpose of comparison, we focused on the estimated signal area, which is the most critical parameter for quantifying metabolites or their isotopic content.

We generated a set of spectra (with varying signal-to-noise ratios ranging from 2.5 to 50) containing two partially overlapping signals: a triplet and a quadruplet, as described in Section 1, separated by 10 Hz (case of partly overlapped peaks, as shown in the bottom spectrum on Figure S2C). The accuracy and precision of the estimated signal areas are presented in Figure S3. At high S/N ratios (>20), both precision and accuracy were comparable when using singlets or multiplets. However, as the S/N decreased, both precision and accuracy improved when multiplets were used. For example, at a S/N ratio of 2.5, the precision (i.e., standard deviation) was 29% for singlets and 13% for multiplets, while the accuracy (i.e., mean error) was 11% for singlets and 2% for multiplets.

Overall, these results demonstrate that explicitly incorporating signal multiplicity into the fitting models enhances the quality of quantification results.

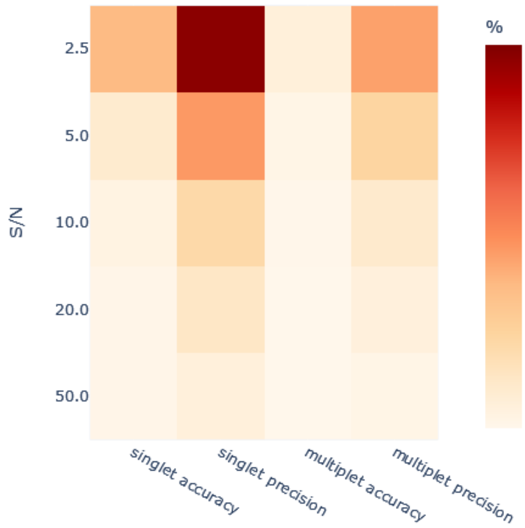


**Figure S3.** Comparison of quantification results obtained by fitting a spectrum with overlapping signals using either singlets or mutliplet.

### 2. Application on metabolomics and isotopic datasets

#### 2.1. Sample preparation

##### 2.1.1. Expression and preparation of enzymes for real-time monitoring of biochemical reactions

To obtain purified glucokinase, glucose-6-phosphate isomerase, and phosphofructokinase A enzymes of *E. coli*, we used the corresponding strains from the ASKA collection (Kitagawa *et al.*, 2006). The cells were grown in 100 mL 2xYT medium supplemented with 30 µg/mL chloramphenicol with vigorous shaking at 37°C to an OD600 ~0.6, then the culture was shifted to 20°C, and isopropyl β-D-1-thiogalactopyranoside was added up to 1 mM to induce expression. After an additional 16 hours of growth, the cells were harvested by centrifugation and lysed by sonication in the buffer containing 50 mM Tris-HCl, pH 8.0, 300 mM NaCl, 5 mM imidazole, 0.5% Triton X100, 2 mM β-mercaptoethanol, and 2 mM phenylmethylsulfonyl fluoride. The cell lysate, clarified by centrifugation at 16,000 x g at 4°C, was applied to a 1-mL HiTrap Talon Crude column (Cytiva). The column was washed with 10 mL of buffer containing 20 mM Tris-HCl, pH 8.0, 300 mM NaCl, 5 mM imidazole, and 2 mM β-mercaptoethanol. Proteins were eluted with 300 mM imidazole, pH 8.0 solution containing 50 mM NaCl and 2 mM β-mercaptoethanol. The collected protein fractions were concentrated and buffer-exchanged to 50 mM Na-phosphate, pH 7.5 containing 50 mM NaCl using Amicon Ultra-4 with 30 kDa cut-off (Millipore), and flash-frozen in liquid nitrogen. The quality of the resulting protein samples was assessed on SDS PAAG. Protein concentration was determined by absorbance at 280 nm using the corresponding calculated extinction coefficients.
The first reaction (for $^{1}H$ & $^{31}P$ experiments) was carried out in a buffer solution (20 mM Na phosphate, 30 mM $MgCl_2$, 50 mM NaCl, 10 mM ATP, and 1 mM TSP (3-(Trimethylsilyl)propionic-2,2,3,3-$d_4$ acid), at pH 7.6) supplemented with 5 mM glucose. Reaction was initiated by addition of glucokinase (0.01 µM), glucose-6-phosphate isomerase (0.45 µM) and phosphofructokinase A (0.27 µM).

The second reaction (for $^{13}C$ experiment) was carried out in a buffer solution (50 mM Na phosphate, 15 mM $MgCl_2$, 50 mM NaCl, 10 mM ATP, and 1 mM TSP (3-(Trimethylsilyl)propionic-2,2,3,3-$d_4$ acid), at pH 7.6) supplemented with 5 mM uniformly $^{13}C$-labeled glucose. Reaction was initiated by addition of glucokinase (0.01 µM).

##### 2.2.2. Cultivation and sample preparation for $^{13}C$-isotope labeling experiments

###### 2.2.2.1. $^{13}C$-labeled acetate

*Escherichia coli* K-12 BW25113 strain was grown in M9 mineral medium complemented with 15 mM glucose (80 % 2-$^{13}C$-D-Glucose / 20 % U-$^{13}C_6$-D-Glucose) and 10 mM 2-$^{13}C_1$-acetate (prepared in solution at pH 7). Cells were grown in flasks at 37°C and 200 rpm, in a volume of 50 mL, as detailed in (Enjalbert *et al.*, 2017). 180 µL samples of medium were filtered (0.2 µm syringe filter, Sartorius, Germany) to remove the cells, and the supernatant was mixed with 20 µL of TSP (10 mM) before analysis.

##### 2.2.2.1. $^{13}C$-labeled aspartate

*Escherichia coli* K-12 BW25113 strain was grown in M9 mineral medium complemented with 60 mM acetate (25 % 12C-acetate, 25 % 1-$^{13}C_1$-acetate, 25 % 2-$^{13}C_1$-acetate, 25 % U-$^{13}C_2$-acetate) prepared in solution at pH 7. Cells were grown in flasks at 37°C and 200 rpm, in a volume of 50 mL, as detailed in Heuillet *et al.* (Heuillet *et al.*, 2018). To obtain proteinogenic amino acids, cells were collected by centrifugation and hydrolyzed at 110°C during 12h in 6N HCl.

### 2.2. NMR spectroscopy and data processing

#### 2.2.1. Real-time analysis of biochemical reactions by $^1H$ and $^{31}P$ NMR

NMR data were acquired at 20ºC on a Bruker AVANCE NEO 800 MHz spectrometer equipped with a 5-mm cryoprobe with z pulsed field gradients. The experiment was carried out in a pseudo-2D manner with simultaneous detection of both $^1H$ and $^{31}P$ nuclei. The following spectral settings for $^1H$ were used: spectral width of 8196.22 Hz corresponding to 10.2442 ppm, acquisition time of 0.999 s, 8 scans per increment and d1 of 4 s. 1024 points were acquired in the indirect dimension (time dimension). Each spectrum is then separated by ns $\times$ (aq +d1) = c.a. 40 s.
The $^1H$ dataset was processed as follow:

- TSP was used for spectrum referencing and normalization, a singlet was fitted on its methyl signal.
- Glucose (Glc) and Glucose-6-Phosphate (G6P) display an apparent triplet in the region of 3.2-3.4 ppm corresponding to the H2 proton of the β isomer of each molecule. The integrals of both signals were extracted by fitting simultaneously two triplets on the corresponding region.
- Fructose-6-phosphate (F6P) and Fructose-1,6-phosphate (FBP) were integrated based on signals that were assigned previously (Cox *et al.*, 2021), using singlet models.
- Adenosine triphosphate (ATP) and Adenosine diphosphate (ADP) were monitored based on their aromatic protons at 8.5 ppm (H7) and 8.2 ppm (H12). These 4 signals were analyzed simultaneously as 4 separate singlets.

The $^{31}P$ dataset acquired simultaneously was fitted for the first 40 time points using MultiNMRFit with a singlet model.

Results obtained for the $^1H$ spectra are shown in Figure 2 of the publication, and examples of results obtained for the $^{31}P$ spectra are shown in Figure S4.

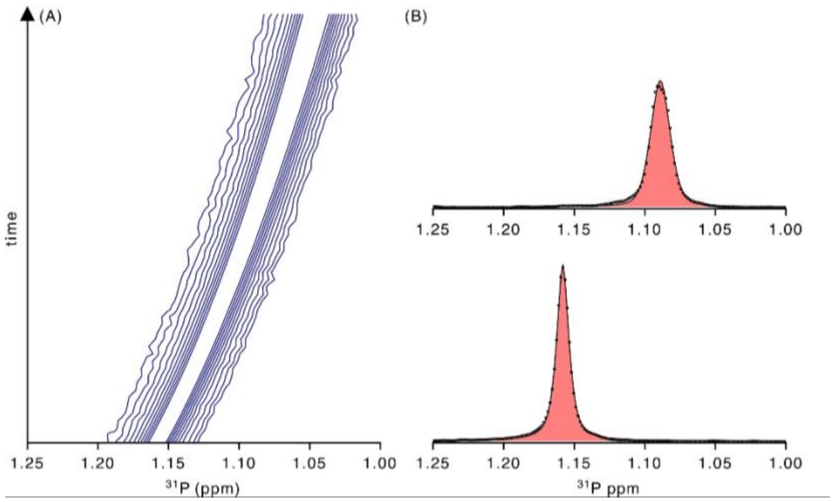


**Figure S4. Example of MultiNMRFit on $^{31}$P data. (A)** Zoom on the pseudo2D $^{31}$P spectra showing the evolution of the signal of inorganic phosphate as a function of time**. (B)** Examples of two $^{31}$P spectra fitted with MultiNMRFit using a singlet, with the experimental and fitted spectra represented by dots and lines respectively,

#### 2.2.2. Real-time analysis of biochemical reactions by $^{13}$C NMR

The $^{13}$C detected experiment was carried out in a pseudo2D manner on a 800 MHz spectrometer. The following spectral settings for $^{13}$C were used: spectral width of 12500 Hz corresponding to 62.1245 ppm, acquisition time of 0.65536 s, 32 scans per increment and d1 of 1 s. 512 points were acquired in the indirect dimension (time dimension). Each spectrum is then separated by ns × (aq +d1) c.a. 53 s.

- Signals of carbon C6 for both anomeric forms of GLC are displayed as doublets of triplets due to the large $^{1}$J(C6-C5) coupling constant and smaller $^{2}$J(C6-C4) and $^{3}$J(C6-C3) (Figure S5A). Triplets were fitted using the triplet model as shown with different colors on Figure S4A for early (left) and later (right) time point.
- Signals of carbons C1 of GLC and G6P have distinct chemical shift upon the anomeric form. The α-C1 carbon resonates near 96 ppm and appear as doublet of triplets due to $^{1}$J(C1-C2), $^{2}$J(C1-C3) and $^{3}$J(C1-C5) and they can all be fitted by triplets (Figure S5B).

The dynamics of the intensity of each signal obtained by MultiNMRFit is shown in Figure S5C.

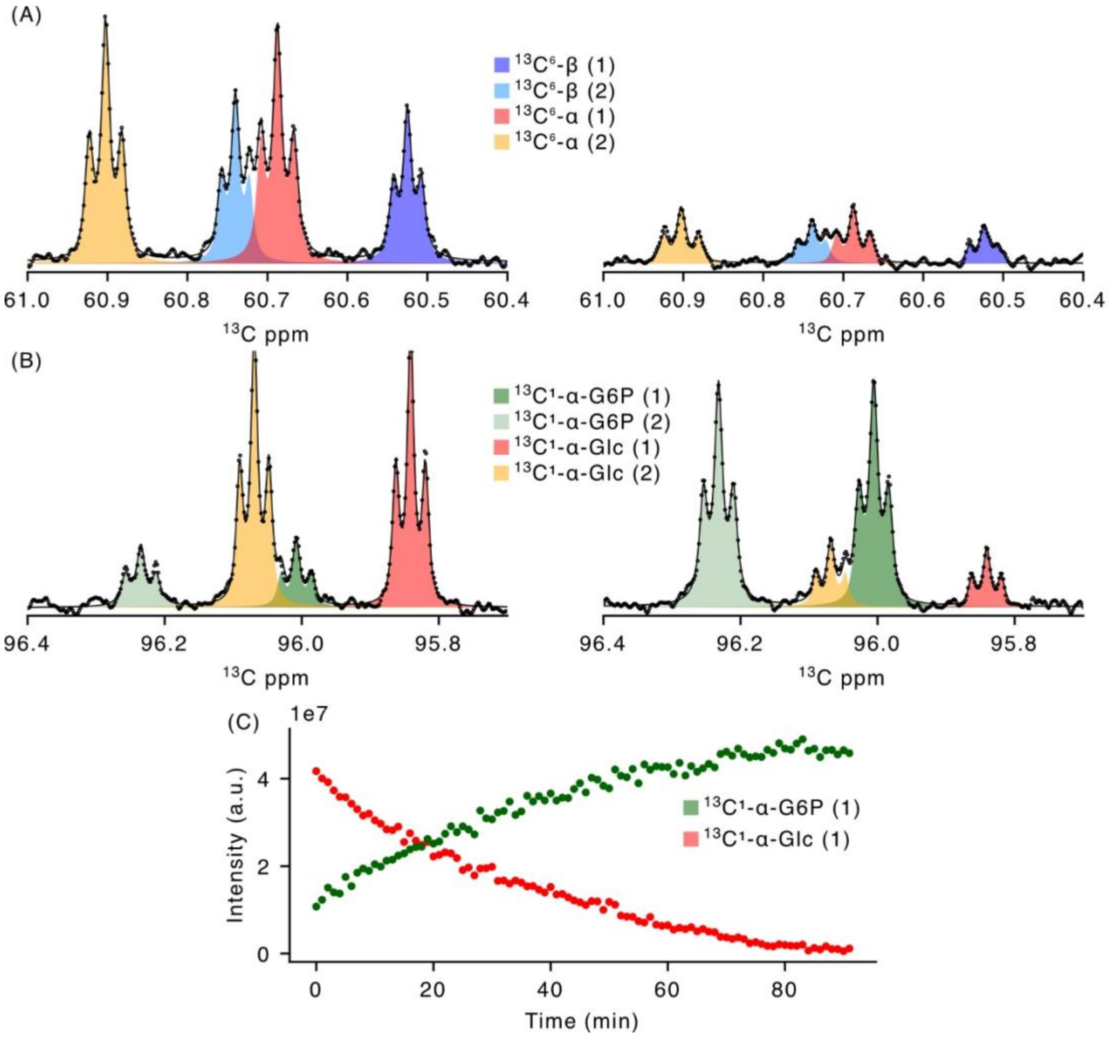


**Figure S5. Application of MultiNMRFit to $^{13}$C NMR spectra. (A)** C6 carbon signals of glucose are shown at two time points for both anomeric forms. **(B)** a-C1 signal of glucose and glucose-6-phopshate. All data were fitted with triplets. For each case individual signals are shown by the shaded areas and the experimental and fitted spectra are represented by dots and lines, respectively. **(C)** Dynamics of integrals for Glc and G6P signals extracted with MultiNMRFit.

#### 2.2.3. Determination of $^{13}$C-isotope incorporation in acetate

Extracellular labeled glucose and labeled acetate concentrations were quantified by 1D $^{1}$H NMR on a Bruker Avance 500 MHz spectrometer equipped with a 5-mm z-gradient BBI probe (Bruker, Germany), as described previously (Millard *et al.*, 2021). Using the following spectral settings for $^{1}$H: spectral width of 10000 (Hz) corresponding to 19.9947 ppm; acquisition time of 3.2767999s, 32 scans per increment and d1 of 3 s. The four isotopic forms of acetate were quantified by fitting the methyl signal at 1.92 ppm using a singlet, two doublets, and a doublet of doublet (Figure S6).

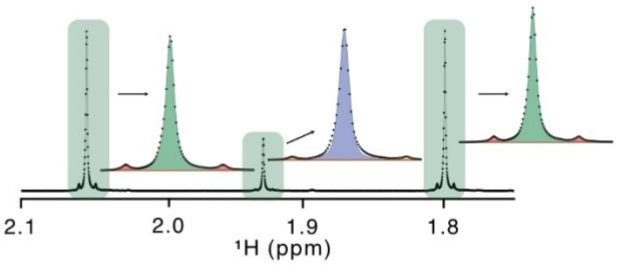


**Figure S6. Experimental spectrum of $^{13}$C-labeled acetate.** Example of spectrum processed with MultiNMRFit with the experimental spectrum shown as dots, the fitted spectrum as a line, and individual signals as shaded areas.

#### 2.2.4. Determination of $^{13}$C-isotope incorporation in aspartate

We used a $^{1}$H-$^{13}$C HSQC experiment to analyse $^{13}$C-incorporation in a standard sample with controlled and known isotopic content, on a Bruker Avance 500 MHz spectrometer equipped with a 5-mm z-gradient BBI probe (Bruker, Germany). Spectral dimensions for the HSQC experiment were 18865 Hz ($F_1$) x 5000 Hz ($F_2$) corresponding to 150 x 9.9974 ppm, with sampling durations of 217 ms ($t_1$), 400 ms (t2). Spectra were centered at 4.700 ppm ($^{1}$H), 75.00 ppm ($^{13}$C). The procedure to extract the 1D $^{13}$C spectra from the 2D $^{1}$H-$^{13}$C was performed according to (Szyperski, 1998). Fitting results are shown in Figure 3D of the publication.
